# Riparian Woody Species Composition, Diversity and Structure in Kiliti Watershed, Northwestern Ethiopia

**DOI:** 10.1101/2025.11.19.689373

**Authors:** Haileyesus Gelaw, Zerihun Woldu, G/medhin Tesefaye, Getahun Haile, Getnet Bitew

## Abstract

Riparian ecosystems are among the most ecologically productive habitats that provide essential ecosystem services, yet they are increasingly threatened by human-induced disturbances. This study examined the composition, diversity, and structural attributes of woody species in the riparian vegetation of the Kiliti watershed, northwestern Ethiopia. 55 systematically distributed sample plots, each 20 × 20 m, were established along riparian corridors. All woody species with diameter at breast height (DBH) ≥ 5 cm were measured for height and diameter, and the number of individuals per species was recorded. Quantitative structural parameters were computed. 95 woody species from 45 families and 75 genera were recorded, dominated by Fabaceae (16%). Shrubs (48%) were the most frequent life form, followed by trees (32%) and lianas (9%). The Shannon–Wiener diversity and evenness values were 3.1 and 0.7, respectively, indicating high diversity and uniform distribution. Diameter and height class distributions exhibited reverse J-shaped patterns, reflecting good regeneration and structural stability. The total basal area (26.28 m^2^/ha) and density values were comparable to other Ethiopian riparian forests. Despite its ecological importance, the Kiliti watershed faces anthropogenic pressure from tree removal. Sustainable management and community-based conservation strategies are vital for maintaining riparian ecosystem integrity.

## 1. Introduction

Riparian ecosystems, situated at the interface between terrestrial and aquatic environments, play a crucial role in maintaining ecological balance, regulating hydrological processes, and supporting high levels of biodiversity (Zaharescu et al., 2017). These zones act as natural buffers (Abood, 2011), filtering sediments and pollutants (Hassanzadeh et al., 2019), stabilizing stream banks (Zaimes et al., 2019), and providing essential habitats for a wide range of plant and animal species (Smith, 2016). In tropical and subtropical regions, riparian vegetation is particularly important for conserving biodiversity and sustaining ecosystem services that local communities rely on for their livelihoods (Davis et al., 2025). However, in many parts of Ethiopia, these fragile ecosystems are increasingly threatened by anthropogenic disturbances such as agricultural expansion, overgrazing, fuelwood collection, and settlement encroachment (Assefa et al., 2020).

Previous studies across Ethiopia have documented significant variations in riparian vegetation composition and structure across different watersheds and climatic zones (Anamo et al., 2023). For instance, research in the Awash and Tana basins revealed that land-use change and watershed degradation have resulted in the loss of native woody species and the dominance of invasive and disturbance-tolerant taxa (Minale and Belete, 2017; Tessema et al., 2020). Other studies have emphasized the ecological importance of riparian corridors for watershed health and biodiversity conservation (Nyemba, 2013; Gebresllassie et al., 2014; Atkinson and Lake, 2020). Despite this growing body of research, information on riparian woody species diversity and structural characteristics in many sub-watersheds, including those in northwestern Ethiopia, remains limited. The Kiliti watershed, located in this region, is one such understudied area where increasing anthropogenic pressures threaten the integrity of riparian habitats (Teshager and Abeje, 2021).

This study seeks to address this knowledge gap by examining the composition, diversity, and structure of riparian woody species in the Kiliti watershed. The central research problem lies in the lack of quantitative data on how land-use dynamics and environmental gradients influence riparian vegetation patterns in the area. Understanding these ecological characteristics is essential for formulating sustainable watershed management and biodiversity conservation strategies (Dinca et al., 2025). Therefore, the specific objectives of this study are to identify and document woody species present in the riparian zones, assess their diversity and structural attributes, and analyze the patterns of species distribution along environmental and disturbance gradients.

The novelty of this study lies in its integrated assessment of species composition, diversity, and structure within a watershed context that has not been previously documented. By combining quantitative ecological data with spatial analysis of riparian vegetation, the research provides baseline information critical for future restoration and management interventions. Moreover, the findings contribute to the broader understanding of riparian ecology in semi-humid highland environments of Ethiopia.

## 2. Material and Methods

### 2.1. Description of the study area

Dangila woreda is situated in the north-western highlands. Dangila town is situated along the Addis Ababa-Bahir Dar road at a distance of 60 km south west of Bahir Dar. The climate is subtropical with annual rainfall around 1600 mm and the main rainy season (known as *Kiremt*) occurring in June–September. The total population of Dangila woreda is estimated at about 200,000 people in an area of about 800 km2. Crop–livestock mixed subsistence farming is the primary source of livelihood. According to a recent survey Belay and Bewket, (2013) some 38 streams and springs flow in the area and 79% of them are used for irrigation. Kiliti Watershed is foundin Dangila woreda, Northwestern part of the country, which is located between 36° 45’ 37.09” E to 36° 49’26” East longitude and 11° 13’ 51” to 11° 16’18” North latitude. The watershed covers a total area of 1759ha (Figure 1). The climatic condition in the watershed is Weyna-dega (midland with 1500-2500m altitude). The altitudinal variation of the watershed area generally ranges from 2072-2343 m.a.s.l. (Teshager and Abeje, 2021). The mean annual rainfall and temperature for the watershed is 2379 mm and 18°C, respectively.

**Figure 1:**
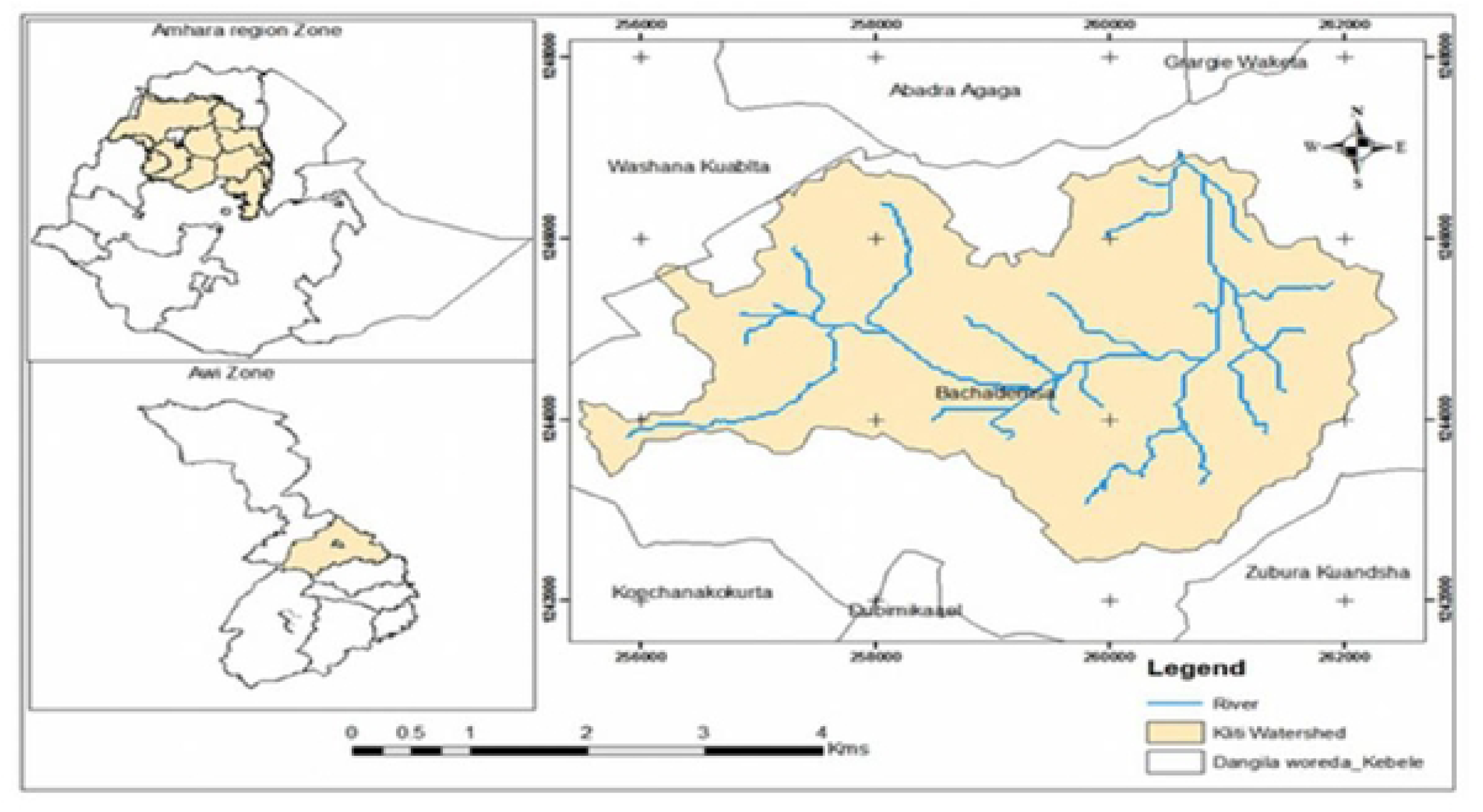
Location map of the study area Awi zone, Dangila wereda, Kiliti watershed. **Source:** Teshager and Abeje (2021).

### 2.2. Sampling design and data collection

A multi-stage sampling procedure were used. In the first stage, within Dangila wereda, Kiliti watershed were selected purposively, due to significant LULC change where occurred over the last three decades (Teshager and Abeje, 2021). Afterward, a preliminary field survey will be conducted to obtain data on the physical conditions of the watershed, including riparian forest size along the streams. In the second stage following the classification by Rosgen (1985), first and second order semiperianal streams such as Amen, Gurdala and Kility were selected. In the third stage, five sample sites were located at various locations of the streams to encompass the range of a priori criteria.

At each sample site three transect lines spaced 500m apart were established (Meragiaw et al., 2018). Transects were laid out preferentially by visual observation of the forest structure of the area (Adugna, 2010). Then, through systematic sampling method within each 500m long transect at a distance of 100m interval 400m2 (20m x 20m) plot were laid (Gemeda et al., 2016; Kebebew and Demissie, 2017; Meragiaw et al., 2018). A total of 55 sample plots were systematically established. Within each plot, all woody plants (trees and shrubs) with a diameter at breast height (DBH) ≥5 cm were measured and recorded. For each individual, the DBH (cm) was measured using a diameter tape at 1.3 m above ground level, and the total height (m) was measured using a hypsometer.

The DBH and height values of all individuals were grouped into six classes: - DBH classes (cm): <5, 5-10, 10-15, 15-20, 20-25, 25-30, >30. Height classes (m): <5, 5-10, 10-15, 15-20, 20-25, 25-30, >30. The frequency of individuals in each class was plotted to examine population structure and regeneration patterns. Each plant was identified to species level using field guides and local vernacular knowledge, and the number of individual stems per species and the number of quadrats (plots) in which each species occurred were recorded. Voucher specimens of unidentified species were collected and later confirmed at the National Herbarium of Ethiopia, Addis Ababa University. Species nomenclature follows Kelbessa and Demissew, 2014.

### 2.3. Data analysis

#### 2.3.1. Diversity

Diversity measures such as Shannon-weiner diversity index (H’), and evenness (E) will be calculated. The formulae for computing diversity indices and evenness will be indicated below:

Shannon-Weiner diversity index (H’)

Shannon-Weiner diversity Index is measured through a combination of species richness (the number of species per sample) and species evenness (the relative abundance of each species). Shannon and weiner (1963) index of diversity will be calculated using the following equation:

H’ = -∑ Pi ln Pi

Where, S = Total number of species

∑ = Summation

Pi = the proportion of individuals or the abundance of the ith species expressed as a proportion of total cover

ln = log base n.

Values of the index usually lie between 1.5 and 3.5 although in exceptional cases, the value can exceed 4.5.

The Shannon’s equitability or evenness (J) of the species was calculated as = H’/ Hmax where Hmax = lnS.

Ln =log basee

S = the number of species

Species richness is the number of species in a given area.

#### 2.3.2. Structural parameters

To describe the structure of riparian woody vegetation, several quantitative parameters were calculated following standard forest ecological methods (Taylor et al., 1993; Ellenberg and Mueller-Dombois, 1974).

**Density** (individuals/ha) was computed as the number of individuals of each species divided by the total sampled area (2.2 ha).

**Basal area** (m^2^/ha) was calculated using the formula: BA = (πd^2^)/4, where d is the DBH. Total basal area per species was then divided by the total sampled area to express dominance in m^2^ per hectare.

**Relative Density (RD)**

(Total number of individual species A/Total number of individuals of all species) X 100

**Relative Dominance** (RDo)

(Dominance of species A/ Dominance of all species) X 100

**Relative Frequency (RF)**

(Frequency of species A/Frequency of all species) X 100

**The Important Value Index** (IVI) for each species was derived as IVI = RD + RDo + RF, which integrates abundance, dominance, and distribution to indicate the overall ecological significance of species within the community.

## 3. Results

### 3.1. Woody species composition and diversity

A total of 95 woody plant species belonging to 75 genera and 45 families was recorded and identified from 55 plots (appendix 1). Fabaceae was the dominant family, with 15 (16%) species. Whereas 26 families with only one species accounted for 27.37% of the total species (Table 1).

**Table 1:**
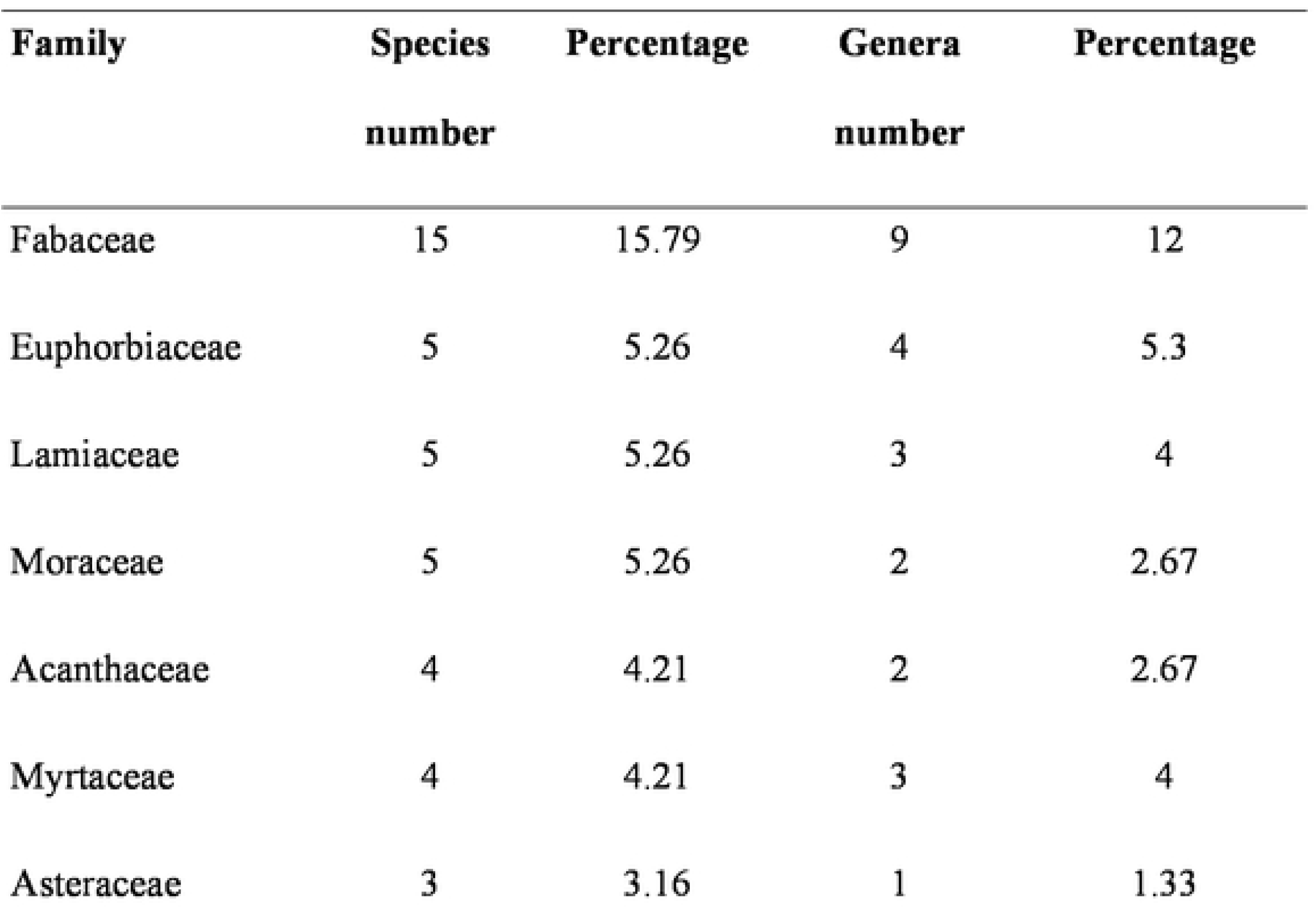
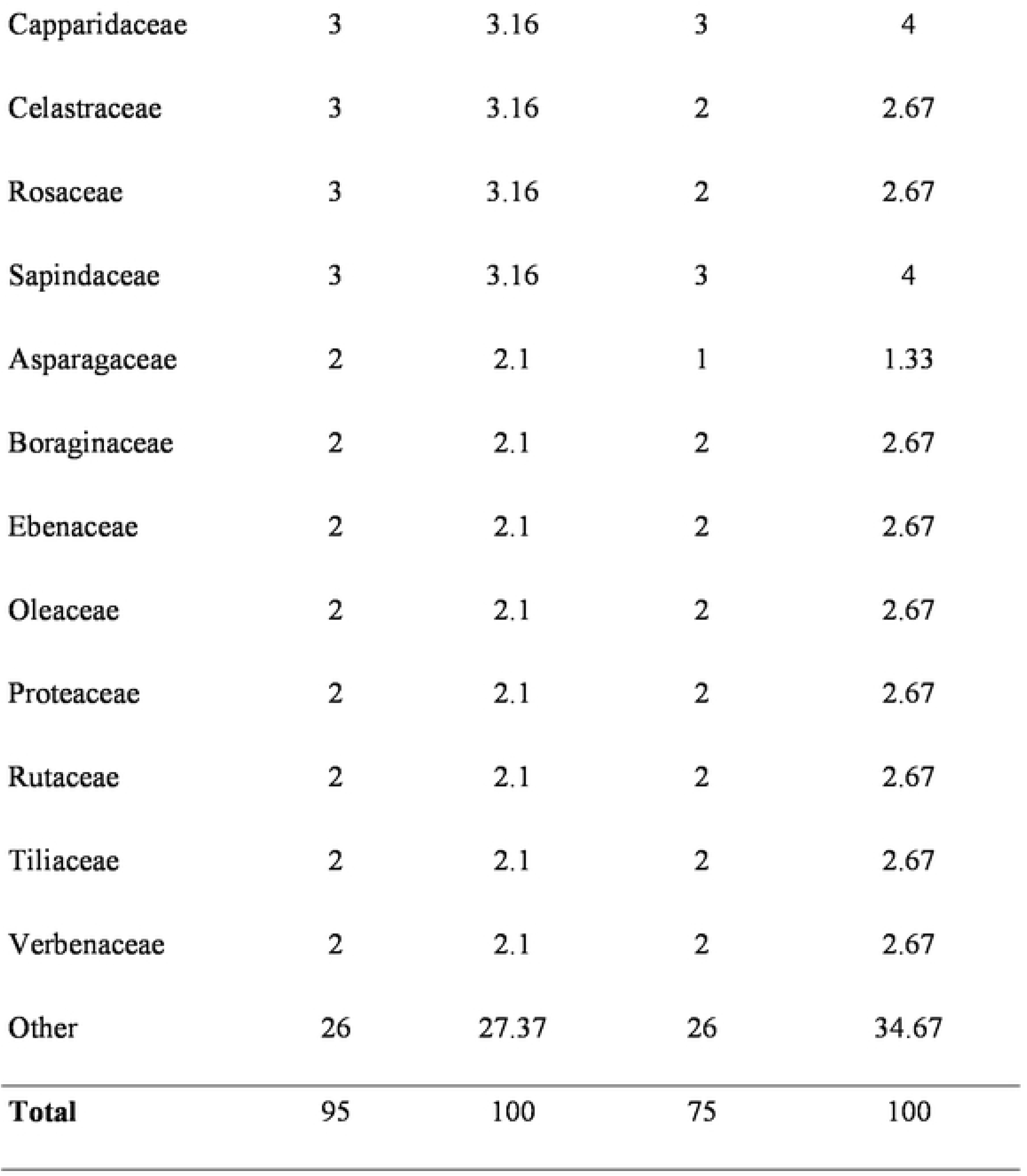
List of families, species, and genera of the riparian woody plant species.

| Family | Species<br>number | Percentage | Genera<br>number | Percentage |
| --- | --- | --- | --- | --- |
| Fabaceae | 15 | 15.79 | 9 | 12 |
| Euphorbiaceae | 5 | 5.26 | 4 | 5.3 |
| Lamiaceae | 5 | 5.26 | 3 | 4 |
| Moraceae | 5 | 5.26 | 2 | 2.67 |
| Acanthaceae | 4 | 4.21 | 2 | 2.67 |
| Myrtaceae | 4 | 4.21 | 3 | 4 |
| Asteraceae | 3 | 3.16 | 1 | 1.33 |
| Capparidaceae | 3 | 3.16 | 3 | 4 |
| Celastraceae | 3 | 3.16 | 2 | 2.67 |
| Rosaceae | 3 | 3.16 | 2 | 2.67 |
| Sapindaceae | 3 | 3.16 | 3 | 4 |
| Asparagaceae | 2 | 2.1 | 1 | 1.33 |
| Boraginaceae | 2 | 2.1 | 2 | 2.67 |
| Ebenaceae | 2 | 2.1 | 2 | 2.67 |
| Oleaceae | 2 | 2.1 | 2 | 2.67 |
| Proteaceae | 2 | 2.1 | 2 | 2.67 |
| Rutaceae | 2 | 2.1 | 2 | 2.67 |
| Tiliaceae | 2 | 2.1 | 2 | 2.67 |
| Verbenaceae | 2 | 2.1 | 2 | 2.67 |
| Other | 26 | 27.37 | 26 | 34.67 |
| **Total** | 95 | 100 | 75 | 100 |

Of the ninety-five woody plant species found in the research watershed, trees accounted for thirty-eight of the species; shrubs and liana accounted for forty-eight and nine species, respectively (Figure. 2). This finding demonstrated that shrubs were adding more species to the total number of woody species found in riparian forest. This indicates that shrubs were quite adaptive and can endure a variety of environmental conditions that impact woody species.

**Figure 2:**
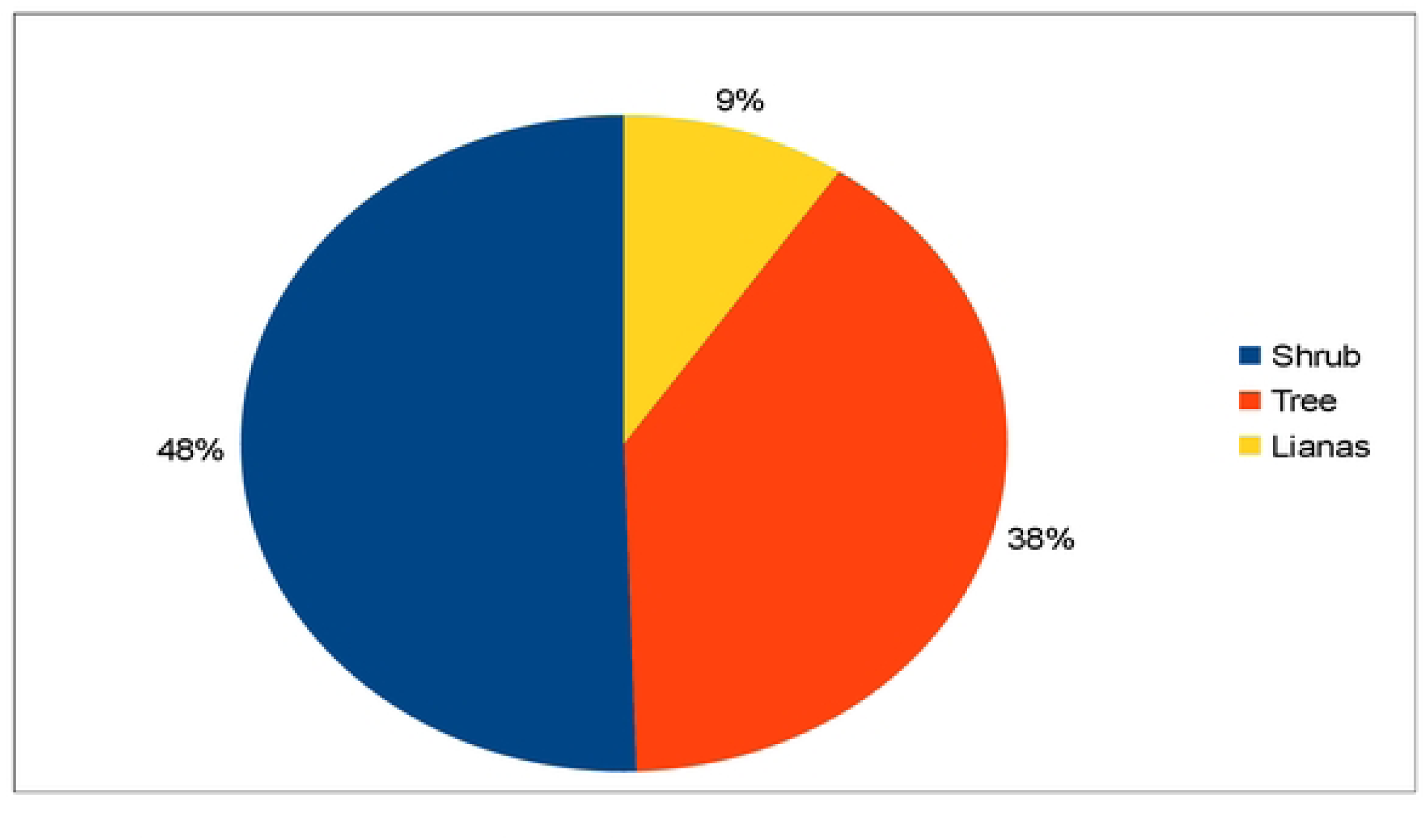
Percentage composition of riparian woody species

### 3.2. Diversity of riparian woody species

In kiliti watershed, the riparian woody plant communities had an evenness value of 0.7 and a Shannon Wiener diversity (H’) of 3.1. This finding indicates that the investigated riparian forest has high species richness, evenness, and Shannon Wiener diversity (Table 2). Riparian forests with the highest evenness values indicate that shrubs species within those forests are uniformly distributed throughout the sampled plots.

**Table 2:** Riparian woody plant Species richness, evenness and diversity.

| **Species**<br>**Richness (S)** | **Diversity index**<br>**(H)** | **True diversity**<br>**(D)** | **Shannon's**<br>**evenness (J)** | **Hmax** |
| --- | --- | --- | --- | --- |
| 95 | 3.1 | 22 | 0.7 | 4.55 |

### 3.3. Density of woody species

The density of riparian woody species in the Kiliti watershed indicate variation among species in terms of stem density per hectare. Some species, particularly *Eucalyptus camaldulensis, Croton macrostachyus, Acacia decurrens and Maytenus arbutifolia* exhibited markedly higher densities compared to others (Figure 3).

**Figure 3:**
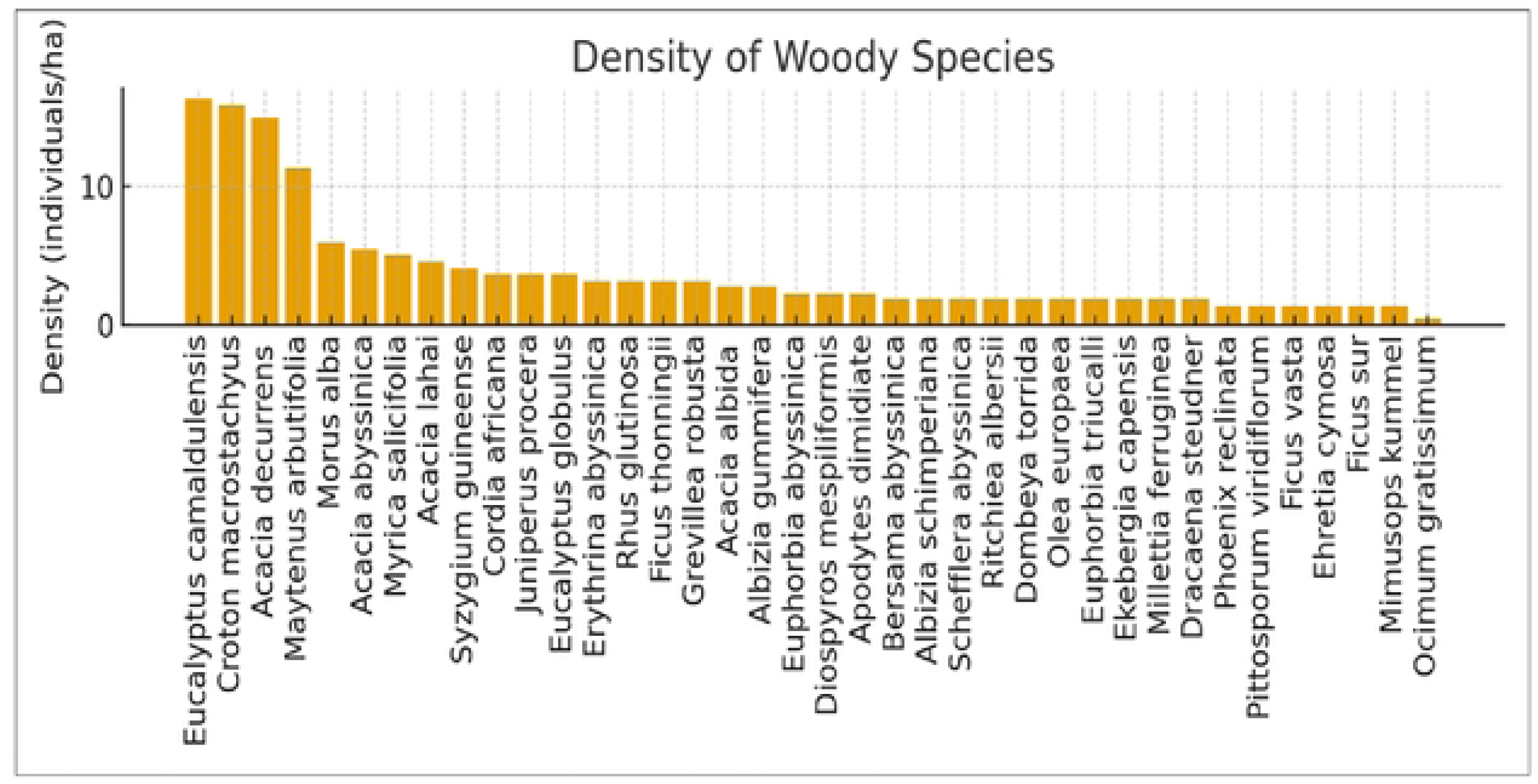
Denisty of woody species (individuals/ha)

### 3.4. Dominance of woody species

Dominance, expressed in terms of basal area contribution per hectare, is shown in Table 3. The results reveal that a few large-sized species such as *Ficus vasta*, *Grevillea robusta*, *Dracaena steudner* and *Ekebergia capensis* contribute disproportionately to the total basal area, while many smaller-stemmed species contribute relatively little. This indicates structural asymmetry within the community, where a small subset of species provides most of the stand’s biomass and canopy coverage.

**Table 3:** Dominance of woody species (Basal area m^2^/ha).

| Scientific name | DO | RDO |
| --- | --- | --- |
| *Ficus vasta* | 5.82 | 22.14 |
| *Grevillea robusta* | 1.66 | 6.31 |
| *Dracaena steudner* | 1.51 | 5.75 |
| *Ekebergia capensis* | 1.5 | 5.69 |
| *Ficus sur* | 1.25 | 4.77 |
| *Ocimum gratissimum* | 1.19 | 4.54 |
| *Albizia gummifera* | 1.08 | 4.12 |
| *Acacia lahai* | 0.98 | 3.73 |
| *Diospyros mespiliformis* | 0.97 | 3.68 |
| Other | 10.32 | 39.26 |
| **Total** | **26.28** | **100** |

### 3.5. Important Value Index (IVI)

The Important Value Index (IVI) of woody species, combining relative density, relative frequency, and relative dominance, is summarized in Table 4. Species with higher IVI values are considered ecologically dominant, either through numerical abundance, wider spatial distribution, or large basal area contribution.

**Table 4:** Important Value Index (IVI) of woody species.

| Scientific name | R.<br>Density | R,<br>Frequency | R.<br>Dominance | IVI |
| --- | --- | --- | --- | --- |
| *Croton macrostachyus* | 10.87 | 22.92 | 0.08 | 33.87 |
| *Acacia decurrens* | 10.25 | 20.83 | 0.79 | 31.87 |
| *Eucalyptus camaldulensis* | 11.18 | 19.44 | 0.16 | 30.78 |
| *Ficus vasta* | 0.93 | 0.69 | 22.14 | 23.77 |
| *Acacia lahai* | 3.11 | 4.17 | 3.73 | 11 |
| *Maytenus arbutifolia* | 7.76 | 2.08 | 0.13 | 9.98 |
| *Acacia abyssinica* | 3.73 | 2.78 | 2.81 | 9.31 |
| *Grevillea robusta* | 2.17 | 0.69 | 6.31 | 9.18 |
| *Ficus sur* | 0.93 | 2.08 | 4.77 | 7.78 |
| Others | 49.07 | 24.31 | 59.08 | 132.46 |
| **Total** | 100 | 100 | 100 | 300 |

### 3.6. Diameter class distribution of woody species

The distribution of woody species across diameter (DBH) classes is illustrated in Figure 4. The DBH histogram reveals that most stems fall within the small to mid-diameter classes, with fewer individuals in larger DBH categories. This general ’reverse-J’ or descending pattern typically indicates ongoing regeneration, where the presence of numerous small stems ensures future replacement of mature trees.

**Figure 4.**
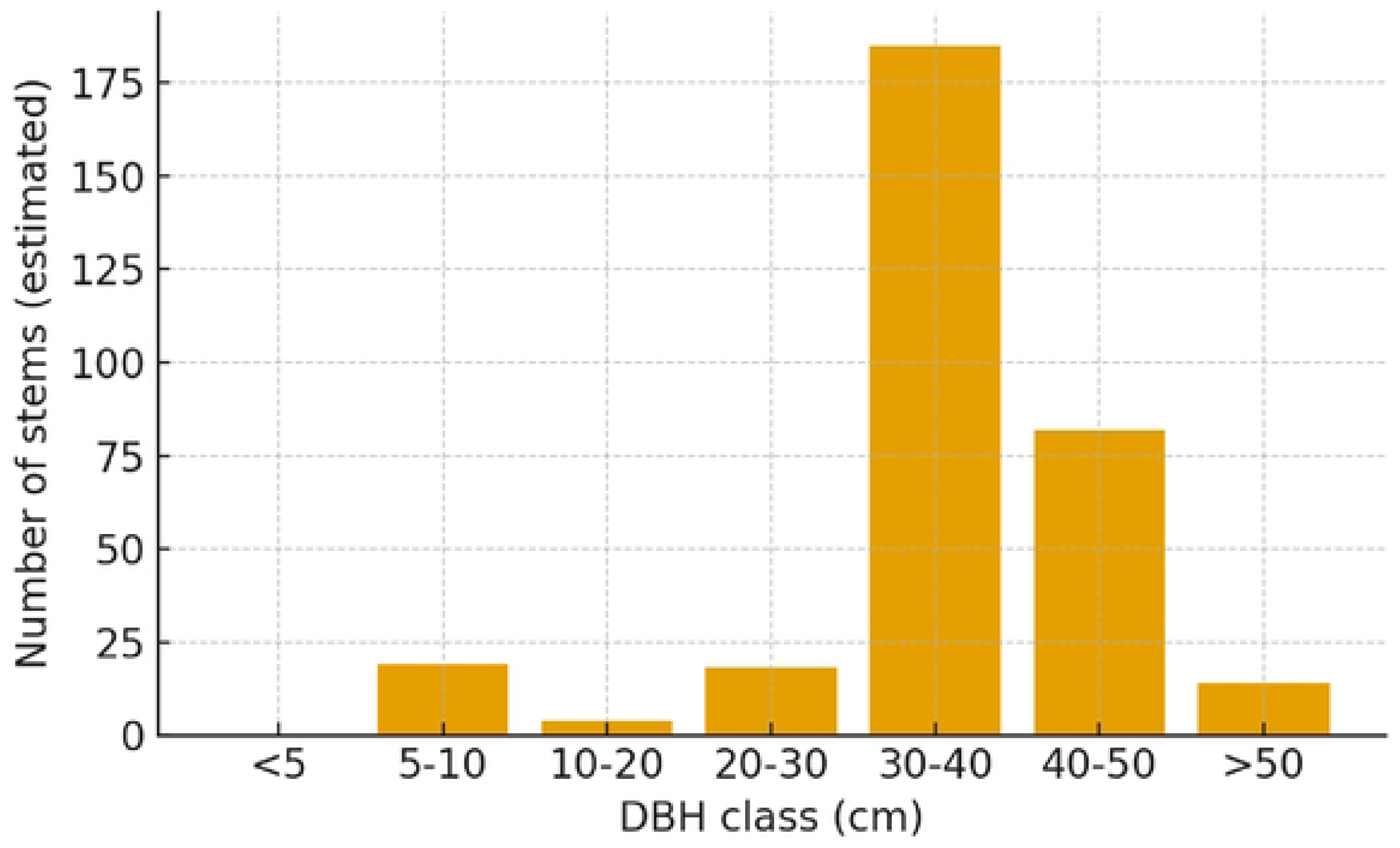
Distribution of woody species across diameter (DBH) classes

### 3.7. Height Class Distribution of Woody Species

Height class distribution of woody species is presented in Figure 5. The majority of individuals occupy lower and intermediate height classes, with relatively fewer trees reaching upper canopy levels. This pattern signifies a mixed-age structure, where younger or sub-canopy individuals dominate in number but larger trees still exist to maintain structural diversity.

**Figure 5:**
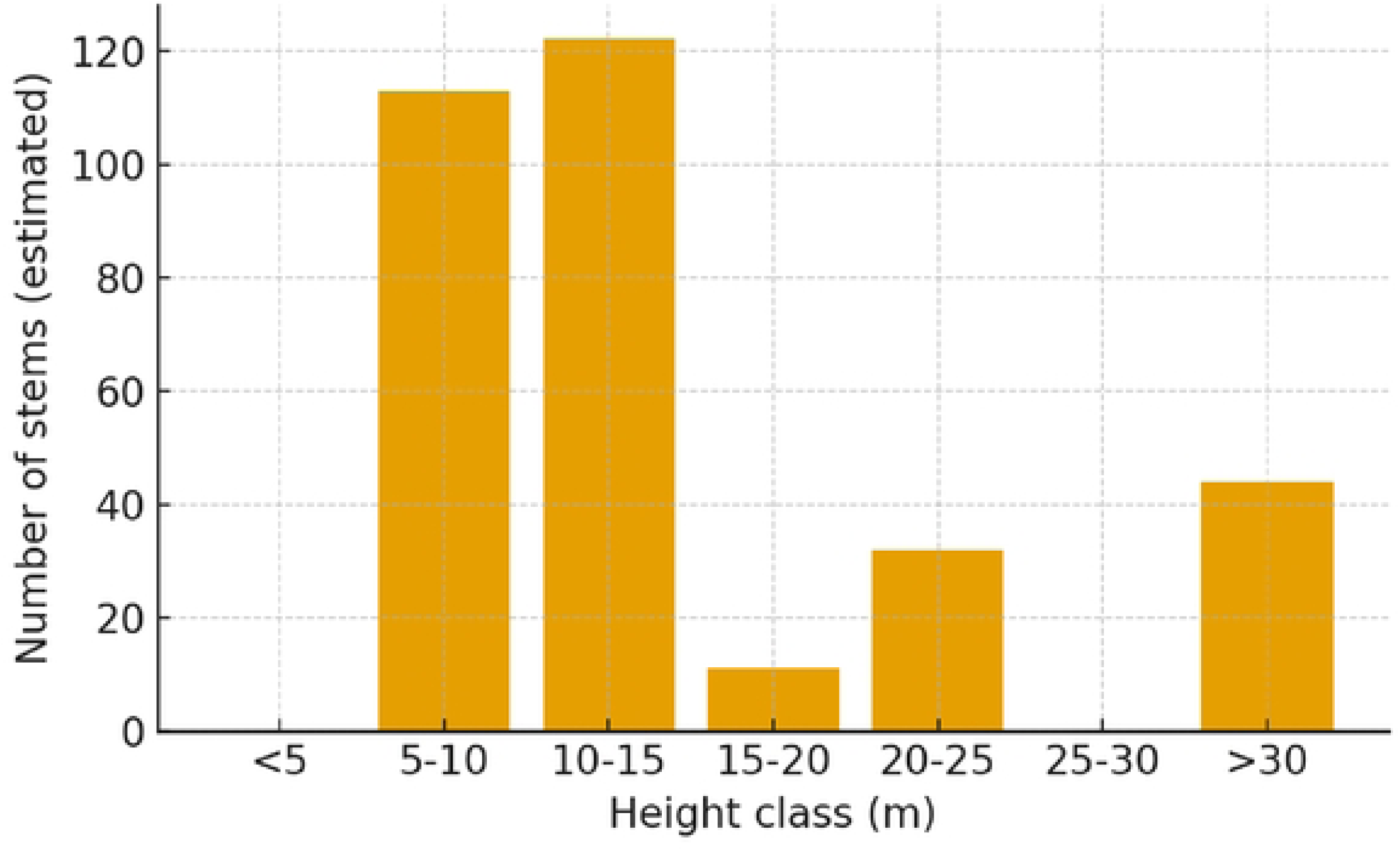
Height class distribution of woody species

### 3.7. Basal area of woody species

The basal area values are shown in Table 5. Basal area serves as a useful measure of stand density and biomass storage. The high contribution from a few species indicates that these taxa dominate the riparian biomass and play a crucial role in carbon storage, soil stabilization, and shading of the riparian corridor.

**Table 5:** Basal area of riparian woody species in kility watershed.

| **Scientific name** | **Basal Area/ha** | **Percentage** |
| --- | --- | --- |
| *Ficus vasta* | 5.82 | 22.15 |
| *Grevillea robusta* | 1.66 | 6.32 |
| *Dracaena steudner* | 1.51 | 5.75 |
| *Ekebergia capensis* | 1.5 | 5.71 |
| *Ficus sur* | 1.25 | 4.76 |
| *Ocimum gratissimum* | 1.19 | 4.53 |
| *Albizia gummifera* | 1.08 | 4.11 |
| *Acacia lahai* | 0.98 | 3.73 |
| *Diospyros mespiliformis* | 0.97 | 3.69 |
| Other | 10.32 | 39.27 |
| **Total** | **26.28** | **100** |

## 4. Discussion

This study revealed that the Fabaceae family holds a dominant position among riparian woody species in the Kiliti watershed. This dominance may result from their efficient pollination strategies, wide ecological amplitude, and successful seed dispersal mechanisms, which allow them to thrive across varied environmental conditions (Tadesse et al., 2017). However, local variations in topography, rainfall, and temperature likely contribute to the observed differences in dominance patterns among plant taxa across sites (Pescador et al., 2015; Masviken et al., 2020).

The Shannon–Wiener diversity index of the Kiliti riparian forest was relatively high compared with other Ethiopian riparian ecosystems, such as Nech Sar National Park (H’ = 2.92) and Dabbis River (H’ = 2.21–3.06) (Mulugeta and Hewan, 2017; Gemeda et al., 2016). This indicates that the Kiliti watershed supports a more diverse woody community, suggesting relatively lower disturbance or better ecological resilience in this ecosystem. The higher diversity could also be attributed to the presence of heterogeneous habitats and the protection offered by local riparian microclimates.

The uneven density distribution of riparian woody species in Kiliti suggests that certain taxa are particularly successful at colonization and persistence, possibly due to differences in environmental tolerance, dispersal strategies, or selective human protection. Similar right-skewed distributions have been documented in other Ethiopian forests, including Ruruki (Belay et al., 2025) and Jibat (Mekonnen et al., 2022), where a few dominant species coexist with numerous rare ones. Such distribution patterns typically characterize secondary or moderately disturbed ecosystems (Hundera et al., 2019).

Species dominance in basal area contribution indicates structural asymmetry, where a small number of species account for most of the stand’s biomass and canopy coverage. Comparable structural patterns have been reported from Ruruki (Belay et al., 2025) and other Afromontane forests (Birhanu et al., 2022; Ayalew et al., 2021). Species with higher Importance Value Index (IVI) scores are often ecologically dominant, reflecting both numerical abundance and wide ecological amplitude. For example, Meragiaw et al. (2018) identified Croton macrostachyus and Syzygium guineense as the most dominant riparian trees along the Walga River, similar to the dominance observed in the Kiliti watershed.

The diameter class distribution of woody species exhibited a characteristic reverse-J pattern, which reflects an abundance of small-sized individuals and a gradual decline toward larger diameter classes. This structure indicates continuous regeneration and recruitment, typical of healthy and self-sustaining forests (Belay et al., 2025; Mekonnen et al., 2023). Likewise, the height-class distribution shows a mixed-age structure where smaller individuals dominate but larger canopy trees persist, maintaining vertical structural diversity. Similar findings were reported by Meragiaw et al. (2018) and Alemu et al. (2020) in other Ethiopian riparian forests.

Overall, the total basal area per hectare in the Kiliti watershed aligns with values reported from other riparian and dry Afromontane forests in Ethiopia (Birhanu et al., 2022; Belay et al., 2025), suggesting a comparable biomass stock and ecological condition. These findings highlight the ecological importance of the Kiliti riparian zone for biodiversity conservation, carbon storage, and hydrological stability.

Despite these promising indicators, the study has limitations. The research was confined to a single watershed and did not include detailed soil, hydrological, or disturbance gradient analyses, which may influence vegetation structure and diversity. Moreover, the study relied on a single-season assessment, potentially overlooking seasonal variations in species phenology and recruitment.

Future research should integrate long-term ecological monitoring, remote sensing, and soil-vegetation interaction studies to better understand the drivers of riparian forest dynamics. Comparative analyses across multiple watersheds and restoration studies would also be valuable to inform sustainable riparian forest management and conservation planning in northwestern Ethiopia.

## 5. Conclusion

The study revealed that the riparian vegetation of the Kiliti watershed supports a diverse assemblage of woody species with varying dominance, density, and distribution patterns. The presence of an inverted J-shaped diameter and height class distribution suggests that the woody vegetation is in a state of active regeneration, dominated by young individuals. However, the reduced representation of larger diameter classes indicates that mature trees are being selectively removed, likely due to human pressure and resource exploitation. The total basal area and overall species density recorded are comparable to those observed in other Ethiopian riparian forests, such as Ruruki and Menagesha Suba, confirming that the Kiliti watershed still retains important ecological value. Nevertheless, anthropogenic disturbances continue to threaten its long-term sustainability. The results emphasize that riparian ecosystems in agricultural landscapes play a crucial role in biodiversity conservation.

## 6. Recommendations

- Strengthen conservation measures: Local authorities and watershed management programs should prioritize riparian zones as key conservation areas and enforce regulations to prevent illegal cutting and encroachment.
- Promote community participation: Awareness creation and participatory management involving local communities are essential to reduce anthropogenic pressure and ensure sustainable utilization of riparian resources.
- Implement restoration interventions: Degraded riparian sections should be rehabilitated through enrichment planting using native species and exclusion of livestock grazing.
- Integrate riparian vegetation management into watershed planning: Riparian buffers should be incorporated into land use plans to enhance ecological connectivity and water quality.
- Future research: Further studies should focus on species regeneration dynamics, soil vegetation relationships, and the effects of human disturbance gradients on riparian ecosystem resilience.

## Acknowledgments

The authors express their sincere gratitude to Injibara University for providing essential facilities and institutional support. We also extend our appreciation to the local community members and field assistants whose cooperation and contributions were invaluable during data collection.

## Data Availability Statement

All relevant data supporting the findings of this study are included in the article’s Supporting Information files.

## Author Contributions

**Conceptualization:** Haileyesus Gelaw, Getahun Haile

**Data curation:** Haileyesus Gelaw

**Formal analysis:** Haileyesus Gelaw

**Investigation:** Haileyesus Gelaw, Getahun Haile

**Methodology:** Haileyesus Gelaw, Zerihun Woldu, Getahun Haile

**Writing – original draft:** Haileyesus Gelaw

**Writing – review & editing:** Zerihun Woldu, Getahun Haile

**Fieldwork:** Getnet Bitew, Getahun Haile

**Supervision:** G/medhin Tesefaye

